# Smell-induced gamma oscillations in human olfactory cortex are required for accurate perception of odor identity

**DOI:** 10.1101/2021.04.15.440045

**Authors:** Qiaohan Yang, Guangyu Zhou, Torben Noto, Jessica Templer, Stephan Schuele, Joshua Rosenow, Gregory Lane, Christina Zelano

**Affiliations:** Department of Neurology, Feinberg School of Medicine. Northwestern University. 320 E Superior Ave, Chicago IL. 60611; Department of Neurosurgery, Feinberg School of Medicine. Northwestern University. 320 E Superior Ave., Chicago IL. 60611

## Abstract

Studies of neuronal oscillations have contributed substantial insight into the mechanisms of visual, auditory and somatosensory perception. However, progress in such research in the human olfactory system has lagged behind. As a result, the electrophysiological properties of the human olfactory system are poorly understood, and in particular, whether stimulus-driven high frequency oscillations play a role in odor processing is unknown. Here, we used direct intracranial recordings from human piriform cortex during an odor identification task to show that three key oscillatory rhythms are an integral part of the human olfactory cortical response to smell: odor induces theta, beta and gamma rhythms in human piriform cortex. We further show that these rhythms have distinct relationships with perceptual behavior. Odor-elicited gamma oscillations occur only during trials in which the odor is accurately perceived, and features of gamma oscillations predict odor identification accuracy, suggesting they are critical for odor identity perception in humans. We also found that the amplitude of high-frequency oscillations is organized by the phase of low frequency signals shortly following sniff onset, only when odor is present. Our findings reinforce previous work on theta oscillations, suggest that gamma oscillations in human piriform cortex are important for perception of odor identity, and constitute a robust identification of the characteristic electrophysiological response to smell in the human brain. Future work will determine whether the distinct oscillations we identified reflect distinct perceptual features of odor stimuli.

## Introduction

Oscillations are ubiquitous across mammalian brain networks [1,2,11,3–10] and studies on their spectrotemporal dynamics have contributed important insight into the mechanisms underlying visual, auditory, and somatosensory perception [10,12–16]. Some of the earliest recordings of brain oscillations occurred in the olfactory system of hedgehogs [17], leading to decades of highly productive animal research on oscillations in the mammalian olfactory system with significant work in rabbit and cat piriform [18–22] and more recently mainly focusing on the rodent olfactory bulb [5,8,31–40,23,41–43,24–30]. However, 80 years later, we still lack understanding of the spectral and temporal dynamics of these oscillations in olfactory processing in the human brain, particularly in higher frequency ranges including beta and gamma.

Gamma oscillations, which comprise the synchronized rhythmic patterns of spiking and synaptic inhibition, have been established as an important mechanism in sensory cortical processing in the mammalian brain [44, 45]. In the visual cortex, gamma oscillations are locally generated and reflect low-level stimulus attributes [46] including grating size, contrast, spatial structure and color [47–50], and are thought to support visual perception by synchronizing the processing and transfer of information within and across areas of visual cortex [51–53]. Similarly, in the auditory cortex, locally generated gamma oscillations may provide a mechanism to integrate neurons according to the similarity of their receptive fields [54], and are involved in pitch perception and sound discrimination [55, 56].

In the olfactory system, odor-driven increases in beta and gamma oscillations have not been consistently identified in direct recordings from human piriform (olfactory) cortex, which has been found to exhibit low frequency oscillations in response to odor [57]. However, in mammals and insects, the cellular and network processes underlying gamma oscillations in the olfactory bulb and cortex have been studied extensively [19,29,40,58–63], and more recent work has begun to provide insight on their functional role as well [23,26,32,33,36,37,40,64], suggesting involvement in organization of sensory information to enable fine odor discrimination and identification [23,65–69]. In addition to gamma oscillations, beta and theta oscillations have been extensively studied in the mammalian olfactory system, and are involved in olfactory learning and respiratory tracking, respectively [18,27,70,71,28,30,33,36,38,40,64,65]. Thus theta, beta and gamma oscillations constitute three key rhythms that have functional significance for olfactory processing in mammals [8]. However, against this background of extensive animal work, the electrophysiological underpinnings of the human olfactory system are vastly understudied. Only a few intracranial studies have directly measured odor-induced oscillations in the human brain; in the amygdala [72–76] and piriform cortex [57]. Human piriform recordings suggest an important role for low frequency (< 8 Hz) oscillations in olfactory processing [57]; theta oscillations were consistently found in piriform cortex within 200 ms of sniff onset during an odor detection task, and their features could be used to decode odor identity. However, the full spectral characteristics of early and later emerging odor responses in human olfactory cortex remain unknown. Furthermore, the functional role of high frequency oscillations in human piriform cortex is virtually unexplored.

In this study, we used stereotactic intracranial electroencephalography (iEEG) to test three main hypotheses about neural responses in human piriform cortex during an olfactory identification task. First, based on the fundamental role of gamma oscillations in human sensory perception and the established rhythms in the rodent olfactory system, we hypothesized that odors would induce theta beta and gamma oscillations in human piriform cortex. Second, based on the fact that olfactory oscillations are modulated by sniffing [26,58,62,63,77–79], which is inextricably linked to odor onset times, we hypothesized that the timing of responses relative to sniff onset would vary across the different frequency bands. Third, based on established links between gamma/beta oscillations and sensory coding [27,30,32,33,35,36,38,64], and the fact that activity in rodent piriform cortex can be used to decode odor identity [80–82], we hypothesized that human piriform beta and gamma oscillations would emerge only during trials where odors were accurately identified, and that features of their oscillatory rhythms would predict odor identification accuracy.

Our results show that odors elicit a stereotyped oscillatory response in human piriform cortex that is consistent in timing and frequency composition across subjects: Immediately following odor onset, early theta increases are quickly followed by gamma and beta increases. The phase of early low frequency responses drives the amplitude of high frequency rhythms only when odor is present. Finally, the distinct oscillatory rhythms we identified were differentially related to behavioral performance, suggesting that these rhythms could potentially give rise to non-interfering representations of different features of odor stimuli [81, 82], though future work is needed to test this hypothesis.

## Results

To examine odor-driven oscillations in human olfactory cortex, we recorded local field potentials from 7 participants who took part in an odor identification task. Each participant’s clinical electrode coverage included piriform cortex (**Fig 1A**). The olfactory task was performed during clinical recording of ongoing electrophysiological activity, at sampling rates ranging between 500 to 2000 Hz, using a 256-channel clinical EEG acquisition system (Nihon Kohden). Each trial began with an auditory cue signaling that odor would be presented, and providing the potential identity of the odor (rose or mint). After a jittered delay (3–7 s), odor was presented to the participant while sniff onset was precisely measured via a pressure sensor at the nose (Salter Labs). On each trial, participants identified the odor by indicating whether or not it matched the identity prompt (**Fig 1B**). To isolate effects driven by odor from those driven by inhalation, which also impacts oscillations in human piriform cortex [78,83–85], we analyzed and directly compared LFPs during inhalations that contained odor and those that did not, resulting in two experimental conditions: odor and no-odor (**Fig 1B**, areas shaded in pink and blue).

**Fig 1.**
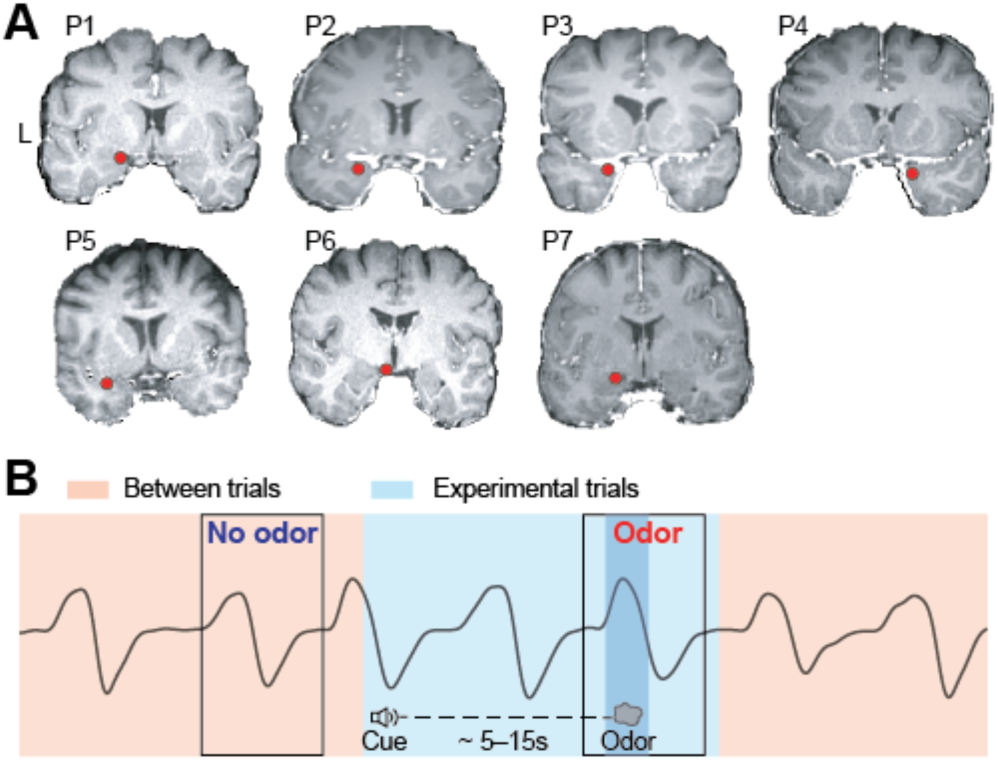
Electrode contact locations and experimental design. (A) The location of the piriform cortex electrode contact (red dot) is shown on each participant’s (P1–P7) brain image. L, left hemisphere. (B) Schematic illustration of the olfactory task, showing odor and no-odor conditions drawn on an illustrative breathing signal (black line). The light blue area indicates experimental trials (starting from cue delivery to the end of the breathing cycle in which odor was delivered), and the pink area indicates between trials. The black boxes indicate the analysis time windows for odor and no-odor trials. The darker blue area indicates odor delivery.

### Odor elicits oscillations in theta, beta and gamma frequency bands in human piriform cortex

To test the hypothesis that odor elicits LFP oscillations in the theta, beta and gamma frequency bands in human piriform cortex, we first conducted a time-frequency analysis combining data from all trials and participants. We computed spectrograms aligned to sniff onset, for odor and no-odor conditions separately. In the odor condition, we found statistically significant increases in theta, beta and gamma frequency bands (**Fig 2A**, left panel) (theta peak: 4.72 Hz, max z score = 14.07; beta peak: 18.98 Hz, max z score = 8.11; gamma peak: 84.94 Hz, max z score = 4.88). In the no-odor condition, we found smaller but significant increases in theta only (**Fig 2A**, middle panel; peak: 4.02 Hz, max z score = 3.44). To quantify the difference between odor and no-odor conditions, we conducted a direct statistical comparison by permuting condition labels to generate a map of z-normalized amplitude differences (**Fig 2A**, right panel). We found that theta, beta and gamma amplitudes were significantly higher in the odor compared to the no-odor condition (permutation test; *P* < 0.05, FDR corrected).

**Fig 2.**
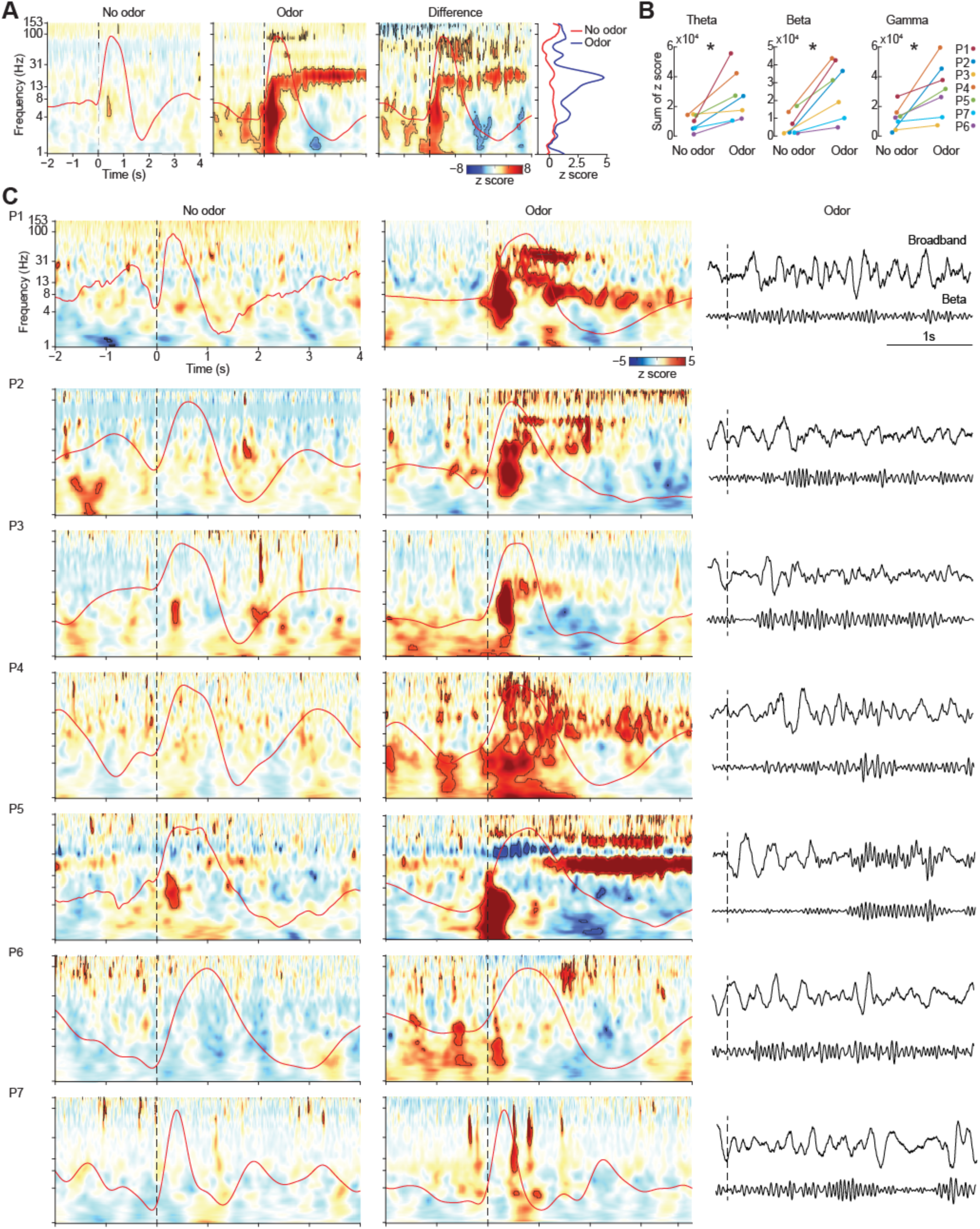
Odor induces theta, beta and gamma oscillations in human piriform cortex. (A) Results from combined analysis showing z-normalized amplitude spectrograms for no-odor and odor conditions (left and middle panels), and the difference between the two conditions (right panel). Average respiratory signal is shown on each panel (overlaid red line). Dashed line represents inhale onset. For odor and no-odor spectrograms, the black contour lines indicate statistical significance of permutation tests against pre-cue baseline (*P* < 0.05, FDR corrected). For the difference spectrogram, the black contour lines indicate statistical significance of permutation of condition labels (*P* < 0.05, FDR corrected). The far-right panel shows average z scores from 0–4 s post sniff across frequencies for odor and no-odor trials. B) Individual theta, beta and gamma summed magnitudes over 0–2 s post sniff. Each dot represents data from a single participant, and lines connect data within the same participant. Asterisk indicates statistical significance (two-tailed paired t test; *P* < 0.05). (C) Z-normalized amplitude spectrograms for each participant for no-odor (left) and odor (middle) conditions, with the average respiratory signal shown on each panel (overlaid red line). Dashed lines represent inhale onset. Black contour lines indicate statistical significance of permutation tests against pre-cue baseline (*P* < 0.05, FDR corrected). Raw broadband and beta (13–30 Hz) time series of a representative trial is shown on the right. The source data for panels (A)–(C) are available in S1 Data.

To confirm our findings at the individual level, we next computed sniff-aligned spectrograms, in each participant separately (**Fig 2B–C**). To quantify responses in each participant, we used standard human EEG frequency band definitions (theta: 4–8 Hz; beta: 13–30 Hz; gamma: > 30 Hz), and looked for responses occurring within these standard frequency ranges in each participant. We computed a direct statistical comparison between the two conditions across participants, which showed that amplitudes in these three frequency bands were significantly stronger across participants during the odor compared to the no-odor condition (**Fig 2B**) (two-tailed paired t test, theta: T_6_ = 3.19, *P* = 0.0187; beta: T_6_ = 4.17, *P* = 0.0059; gamma: T_6_ = 2.98, *P* = 0.0245). Notably, in every single participant, theta, beta and gamma amplitudes were larger during the odor compared to the no-odor condition (**Fig 2B**, each dot is the result from one participant and lines connect same-participant results between conditions). Though there was some variation across individuals in the frequency of responses, these effects were evident in the individual spectrograms of most participants (**Fig 2C;** black outlines indicate statistical significance via permutation test; *P* < 0.05, FDR corrected), and in the minimally-processed signals shown next to each individual spectrogram (**Fig 2C**, right). In each participant, theta, beta and gamma responses were maximal inside piriform cortex, with significantly reduced responses outside of piriform on the same depth wire (two-tailed paired t test; theta: *P* = 0.0045, T_6_ = 4.42; beta: *P* = 0.014, T_6_ = 3.44; gamma: *P* = 0.0070, T_6_ = 4.02) (**S1 Text**; **S1 Fig 3A**). To further visualize the spatial distribution of responses in each frequency band, we created heat maps of responses across all implanted electrodes in all participants, projected onto an axial slice through piriform cortex (**S1 Fig 3B**). This allowed visualization of response magnitudes in the anterior-posterior direction, with the superior-inferior axis flattened. A hot spot of increased activity can be seen corresponding to piriform cortex (**S1 Fig 3B**, pink boxed areas; axial slice contains piriform cortex).

To address the possibility that the responses we observed were due to the steep slopes of sensory-evoked potentials, we conducted an analysis to separate the spectrograms into odor-evoked and odor-induced effects [86–88]. In this analysis, the odor-evoked spectrogram includes only the phase-locked power, which contributes to the sensory-evoked potential, whereas the odor-induced spectrogram includes only the non-phase-locked power, which is oscillatory and is not related to the ERP [89, 90] (see methods). Results of this analysis showed that odor-induced oscillations were not eliminated when the phase-locked components of the signal were removed (**S1 Fig 1**) and thus were not due to the steep slope of sensory-evoked potentials.

In rodents, odor has been shown to elicit responses in multiple gamma subbands [8,64,91]. To examine gamma effects more closely and to look for multiple gamma sub-bands in humans, we re-computed individual spectrograms using a linear, as opposed to log, frequency scale. We found that most subjects showed odor-induced oscillations right around or just below 30 Hz. Most subjects also showed a higher frequency broadband response, around 90–150 Hz (**S1 Fig 2**).

Together, these results suggest that odor consistently induces distinct theta, beta and gamma oscillations in human piriform cortex.

### The time course of olfactory cortical oscillations varies across frequency bands

The spectrograms in **Fig 2** showed apparent variable time courses of oscillations relative to sniff onset across frequency bands (**Figs 2A and 2B**). Theta appeared to emerge and dissipate soonest, with beta and gamma emerging later and persisting longer. To quantify these differences, we conducted a series of analyses to characterize the timing of responses across frequency bands. These included a percent-change analysis to examine oscillatory increases over time, a bootstrapping analysis to minimize potential impact of noisy trials, a cluster-based analysis to quantify the exact timing and magnitude of continuously significant increases in oscillatory amplitude, and a circular-distribution analysis to examine oscillatory peaks over respiratory phase.

In our first analysis, we calculated the percent change in amplitude for each frequency band at each time point, for each condition (**Fig 3A**). This allowed us to determine the timing of the emergence of significant differences between odor and no-odor conditions (two-tailed paired t test; *P* < 0.05, FDR corrected). This analysis showed distinct temporal dynamics of odor responses for each frequency (**Fig 3A**, black lines above each panel). Theta oscillations were the earliest to emerge and dissipate, beginning 126 ms prior to sniff onset and ending 516 ms after sniff onset. Gamma oscillations emerged next, 116 ms after sniff onset, intermittently persisted through exhalation, and dissipated 3678 ms after sniff onset. Beta oscillations emerged last, 144 ms after sniff onset, and persisted longest, ending 3786 ms after sniff onset.

**Fig 3.**
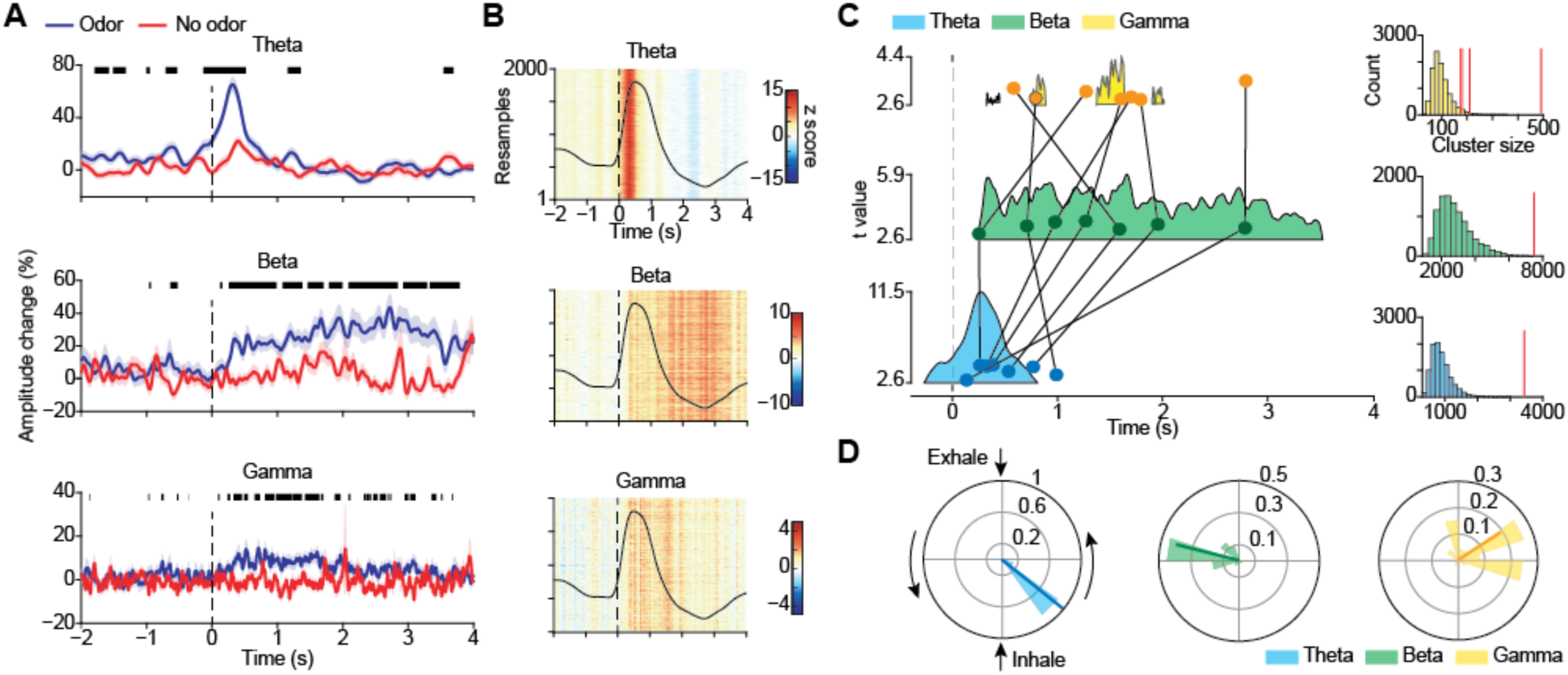
Odor-induced theta, beta and gamma oscillations have different time courses relative to sniff onset. (A) Percent-amplitude-change time series for odor (blue line) and no-odor (red line) conditions for each frequency band. Shaded areas indicate one standard deviation from the mean across all trials. Thick black lines above indicate statistically significant differences between conditions (FDR corrected *P* < 0.05, paired t test). (B) Bootstrap colormap of z-normalized amplitude for each frequency band, with results from each repetition stacked along the Y axis. Vertical stripes of consistent color indicate consistent amplitudes across subsampled sets of trials. (C) Colored shaded areas represent clusters of statistical significance of the trial-by-trial baseline-corrected amplitude time series, for each frequency band (cluster correction test *P* < 0.05, corrected). Upper boundaries of the shaded areas represent the t statistic from the one-sample t test of baseline-corrected amplitude at each time point against 0. Scales for t statistics are shown to the left of each cluster. Overlaid darker dots indicate the average timing of the bootstrapped distribution of peak amplitude occurrence for each frequency band, for each participant. Each dot represents data from one participant. Dots from the same participant are connected by black lines. On the right, cluster masses in relation to the permuted null distribution for each frequency band are shown, with actual values represented by the vertical red line. (D) Respiratory phase angle distributions corresponding to bootstrapped peak amplitudes for each frequency band. Scale of the polar histogram represents pdf of the distribution. Darker lines overlaid on the distribution indicate the mean vector of the phase distribution. Angle and radius represent averaged phase angle and PLV value respectively. The source data for panels (A)–(D) are available S2 Data.

In a second analysis, designed to minimize the contribution of noisy trials and to statistically evaluate the time courses, we tested whether differences in temporal dynamics of responses across frequency bands were stable across resampled sets of trials. We conducted a bootstrapping analysis to create separate profiles of the timing of emergence of responses for each frequency band. We created a z-normalized time series of resampled mean values for each frequency band, represented as a colormap (**Fig 3B**, y-axis is each bootstrap repetition). We found stable time courses across repetitions, indicated by stripes of increased amplitude at particular times for each frequency band. To quantify the timing of amplitude increases, we performed a cluster-based statistical analysis; for each frequency band, we generated a t-statistic at each time point for all z-normalized odor trials, and then conducted a permutation-based analysis to correct for cluster size. We found clusters that corresponded well with the spectrograms shown in Fig 2, with the percent-change analysis, and with the bootstrap analysis (**Fig 3C**). We found one significant cluster for the theta band extending 266 ms pre sniff to 810 ms post sniff (permutation test; *P* < 0.0001), one for the beta band extending from 238 ms to 3524 ms post sniff (*P* < 0.0001), and four separate clusters for the gamma band (326 ms to 452 ms post sniff, *P* = 0.0346; 764 ms to 892 ms post sniff, *P* = 0.0145; 1374 ms to 1642 ms post sniff, *P* < 0.0001; 1902 ms to 2016 ms post sniff, *P* = 0.0451). We subsequently conducted the same bootstrapping analysis at the single subject level, and plotted the average time point of each subject’s bootstrapped peak distribution on top of the significant clusters in Fig 3C (see overlayed dots, each representing the peak value for a single participant, with lines connecting each participant’s values across frequencies). Although single participant data was more variable, beta and gamma peaks occurred later than theta peaks for almost all subjects. These results suggest that the unique time courses of odor responses across frequencies were unlikely to be caused by noise, spikes, and other artifacts.

In a third analysis, designed to account for individual respiratory differences in the time domain, we examined the circular distributions of oscillatory peaks over the respiratory phase. On each repetition of a bootstrapping analysis, the peak amplitude and corresponding phase angle were calculated from the average LFP and respiratory time series. We found that theta peaks narrowly aggregated at early stages of inhale (mean phase angle = –0.67 rad, PLV = 0.99, Rayleigh’s z = 998.05, *P* < 0.0001) (**Fig 3D**, left), consistent with previous findings in humans [57]. By contrast, beta peaks consistently aggregated at the exhale trough (mean phase angle = 2.98 rad, PLV = 0.80, Rayleigh’s z = 647.78, *P* < 0.0001) (**Fig 3D**, middle). Gamma peaks were more broadly distributed, with most centered around inhale peak or the transition between inhale and exhale (mean phase angle = 0.57 rad, PLV = 0.63, Rayleigh’s z = 401.69, *P* < 0.0001) (**Fig 3D**, right). Combined, these findings suggest that following presentation of odor, theta oscillations increase earlier than beta and gamma oscillations, and that while theta and gamma oscillations are maximal during inhalation, beta oscillations peak during exhale.

### Gamma and beta oscillations are required for accurate odor identification

Piriform cortex has been postulated to be a major driver of gamma oscillations in the brains of rodents [18,92] and cats [19–22,93]. Furthermore, numerous rodent studies suggest that beta and gamma oscillations relate to odor learning and discrimination, through both local odor coding [23,33,35,40,64,94] and recruitment of larger-scale networks [27,30,35–37,64]. We therefore hypothesized that beta and gamma oscillations in human piriform cortex would correlate more strongly with task performance than theta. We conducted three separate analyses to explore the relationship between odor-driven LFP oscillations and odor identification accuracy. These included a time-frequency analysis to look for gross differences between correct and incorrect trials, a correlation analysis to look for a relationship between accuracy and oscillations across trials, and a machine learning analysis to determine whether amplitudes of different frequency responses could predict accuracy.

First, in a combined time-frequency analysis, we computed odor-onset aligned spectrograms during correct and incorrect trials separately, in order to look for differences across frequency bands. We found that during trials in which the participant correctly identified the odor, there were statistically significant increases in theta, beta and gamma band amplitudes (**Fig 4A**, left panel; *P* < 0.05, FDR corrected). By contrast, during trials in which the participant failed to identify the odor, statistically significant increases were found only in the theta band, with no such increases in the beta or gamma bands (**Fig 4A**, right panel; *P* < 0.05, FDR corrected; **Fig 4B**, upper panels and lower left panel show the distribution of single trial theta, beta and gamma values for correct and incorrect trials). Since most participants performed above chance on the task, there were a larger number of correct trials than incorrect trials (252 versus 71). To account for this difference, we conducted a resampling analysis with 200 repetitions in which correct and incorrect trials were resampled with replacement to include the same number of trials on each repetition, and we computed the distribution of the differences in amplitude (correct – incorrect) within each frequency band (**Fig 4B**, lower left panel). We found statistically significantly larger responses during correct compared to incorrect trials for beta and gamma bands only, with no significant differences in the theta band. (One-sample t test against 0 for the difference between correct and incorrect trials: theta: T_199_ = 1.21, *P* = 0.2247; beta: T_199_ = 35.55, *P* < 0.0001; gamma: T_199_ = 42.61, *P* < 0.0001. Paired sample t tests for differences between frequency bands: theta vs beta: T_199_ = –41.02, *P* < 0.0001; beta vs gamma: T_199_ = 17.36, *P* < 0.0001; gamma vs theta: T_199_ = 37.10, *P* < 0.0001.). This combined resampling analysis was designed to account for differences in numbers of correct and incorrect trials, which had the advantages of ensuring a balanced comparison across conditions and a robust number of incorrect trials. However, the contribution of individual subjects to the observed effects was unclear. To account for this, we quantified theta, beta and gamma responses for correct and incorrect trials separately, in each participant. Since some participants had no, or just a few, incorrect trials, we limited this analysis to include participants with more than 3 trials of each condition (resulting N = 5). Though numbers of incorrect trials were relatively low for some subjects, results of this analysis confirmed the combined findings: responses in all three frequencies were significantly above zero for correct trials (theta: T_4_ = 4.12, *P* = 0.014 ; beta: T_4_ = 2.87, *P* = 0.045; gamma: T_4_ = 4.71, *P* = 0.0092), whereas only responses in theta band were significantly above zero for incorrect trials (theta: T_4_ = 7.28, *P* = 0.0019; beta: T_4_ = 1.77, *P* = 0.15; gamma: T_4_ = 1.41, *P* = 0.23) (**Fig 4B**, lower right panels). These findings were also apparent in individual participant spectrograms (**S4 Fig 4B**).

**Fig 4.**
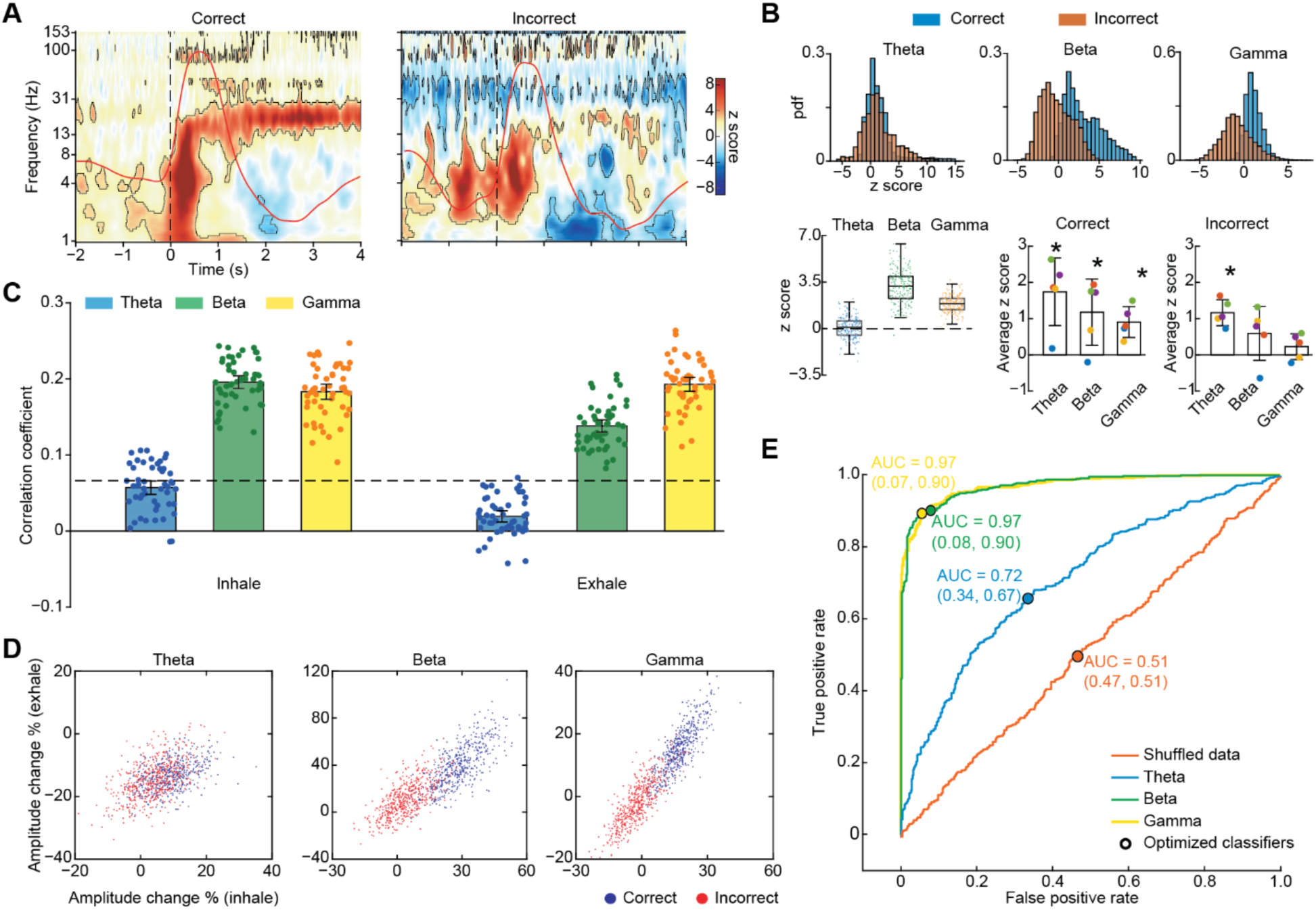
Beta and gamma oscillations predict odor identification accuracy. (A) Z-normalized spectrograms for correct (left) and incorrect (right) trials separately. Averaged respiratory signals are overlayed on each spectrogram (red lines). Dashed lines represent sniff onset. Black contours indicate statistical significance of permutation test against pre-cue baseline (*P* < 0.05, FDR corrected). (B) Amplitudes within each frequency band, for correct and incorrect trials separately. The upper panels show the distribution of z score amplitude values for each trial for the theta, beta and gamma bands, for correct (blue) and incorrect (red) trials. The lower left panel shows box plots for the bootstrapped distributions of the difference between correct and incorrect trials. Boxes represent the 25^th^ to 75^th^ percentile of each distribution, the central marker indicates median, and whiskers extend to the extremes of the data excluding outliers. Colored dots represent raw difference values for each distribution. The lower right panels show individual participant’s mean response amplitudes in the theta, beta and gamma bands, for correct and incorrect trials separately. Each dot represents data from a single participant, bars indicate the mean across participants and error bars are standard error across participants. Stars indicate statistical significance (paired, two-tailed t tests). See Fig 2B for the color code of single participant dots. (C) Bar plots of bootstrapped Pearson’s correlation coefficients computed between oscillatory amplitudes and task accuracy for each frequency band, over the inhale and exhale time periods separately. Error bars indicate 95% confidence interval of the mean. Dashed line indicates FDR threshold for significant r value. Dots represent single correlation coefficient calculated from each bootstrap. (D) Scatter plots show bootstrapped percent change relative to baseline for correct and incorrect trials, during inhale and exhale time periods for each frequency band. Each dot is the average result from one bootstrap repetition. (E) Receiver operating characteristic (ROC) curves of linear binary SVM classifier applied to each of the scatter plots in panel C. Black dots represent optimized classifier, area under the curve (AUC) is indicated for each ROC curve. The source data for panels (A)–(E) are available in S3 Data.

Second, we looked for a relationship between task performance and responses in each frequency band on a trial-by-trial basis. To account for differences within frequency bands over time, we conducted this analysis during the inhale and exhale periods separately. Trials were resampled to generate a distribution of correlation coefficients representing the relationship between response amplitudes and behavioral accuracy for each frequency band across the resampled trial sets. Significant correlations between accuracy and amplitude were evident in the beta and gamma bands during both inhale and exhale (50 bootstraps, t test against 0, FDR corrected threshold for significant correlation coefficient; beta during inhale: T_49_ = 31.25, *P* < 0.0001; beta during exhale: T_49_ = 17.64, *P* < 0.0001; gamma during inhale: T_49_ = 23.70, *P* < 0.0001; gamma during exhale: T_49_ = 28.25, *P* < 0.0001.), but no such correlation was found in the theta band (theta during inhale: T_49_ = –2.04, *P* = 0.97; theta during exhale: T_49_ = –12.90, *P* = 1). In line with this, a direct statistical comparison across frequency bands showed that correlations were significantly stronger between behavioral accuracy and amplitude for beta and gamma bands compared to theta band during both inhale and exhale time periods (**Fig 4C**) (two way repeated measures ANOVA: main effect of time window: F_1, 49_ = 111.08, *P* < 0.0001; main effect of frequency band: F_2, 98_ = 547.98, *P* < 0.0001. Paired sample t test across frequency bands during inhale for theta vs beta: T_49_ = – 29.37, *P* < 0.0001; for beta vs gamma: T_49_ = 3.11, *P* = 0.0031; for gamma vs theta: T_49_ = 24.95, *P* < 0.0001; paired sample t test across frequency bands during exhale for theta vs beta: T_49_ = –26.51, *P* < 0.0001; for beta vs gamma: T_49_ = –11.13, *P* < 0.0001; for gamma vs theta: T_49_ = 33.95, *P* < 0.0001), also see **S4 Fig 4A**.

Finally, we used machine learning to perform a classification analysis on inhale and exhale amplitudes for each frequency band, to determine whether these features could predict task accuracy. To this end, we applied a binary linear support vector machine (SVM) classifier to the bootstrapped data, plotted the receiver operative characteristic (ROC) curves and calculated the area under curve (AUC), which was used together with optimized classifier accuracy as the predictor of data separability. We found that beta and gamma amplitudes were highly successful at separating the trials by accuracy, while theta amplitudes were less so (**Fig 4D**). Specifically, gamma accuracy was maximal at 91.5%, beta accuracy was also high at 91%, whereas theta accuracy was lower, at 66.5%. This same procedure was applied to shuffled data, which resulted in chance performance (accuracy = 52%), validating our methods. These findings correspond well with the results from the two previous analyses, and, taken together, suggest that beta and gamma band amplitudes are more strongly correlated with task performance than theta amplitudes.

### Odor-elicited high-frequency oscillations are not driven by attention

Because the odor condition occurred during experimental trials and the no-odor condition occurred between experimental trials, responses during the odor condition could potentially have been impacted by attentional states that were not present during the no-odor condition. Previous human and animal work suggests that olfactory cortical responses are modulated by attention [95, 96], particularly in lower frequency LFPs [97, 98]. To control for potential effects of attention on our findings, we analyzed odorless sniffs taken in attended states during experimental trials. Each trial began with a cue, followed by a delay (3–7s) prior to presentation of odor. Therefore, during many trials, sniffs of clean air occurred between the cue and odor (**Fig 5A**). These sniffs occurred during attended states, but did not contain odor, thus providing a means to isolate the effects of attention from the effects of odor. Because a sniff did not always occur between the cue and the odor, the number of trials in the attended odorless condition was lower than the odor condition. Therefore, we resampled odor trials to match the number across conditions. We computed spectrograms aligned to attended odorless sniffs and compared them to those aligned to odor sniffs (**Fig 5B**, left and middle). We found that attended odorless sniffs induced oscillations in low frequencies (below 8 Hz) only (*P* < 0.05, FDR corrected), and found no stable higher frequency oscillations (> 8Hz). We then computed the z-normalized difference across conditions (**Fig 5B**, right) and found significant differences between attended odor trials and attended no-odor trials in the theta, beta and gamma bands, such that oscillations in all three frequency bands were increased when odor was present, even after removing the effects of attention (**Fig 5B**, right; *P* < 0.05, FDR corrected). Since theta oscillations were still detected in the attended no-odor condition, these findings suggest that the increases in beta and gamma bands were independent of attention, whereas theta band increases reflected, at least to some degree, attentional states.

**Fig 5.**
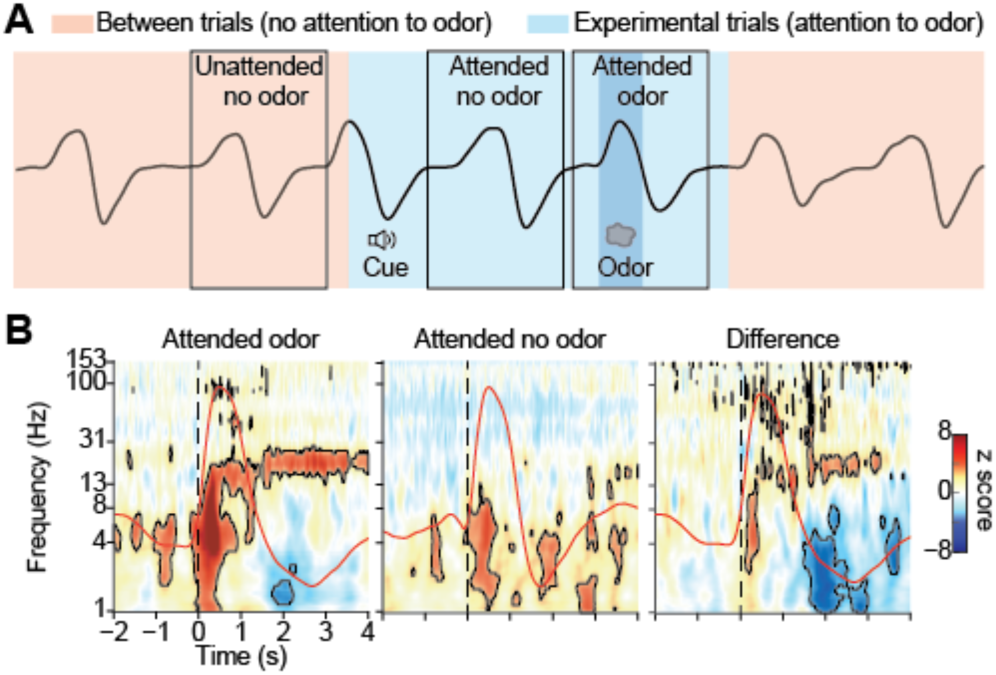
High frequency odor-elicited oscillations are not driven by attention. (A) Schematic illustration of attended and unattended conditions. Attended no-odor trials were defined as breaths taken during experimental trials, between the cue and the odor. No odor was present, but the participant was in an attended state. Blue area indicates the time of experimental trials, and pink areas indicates the time between trials. Black boxes indicate the analysis time windows for the different conditions. (B) Z-normalized spectrograms of attended odor and attended no-odor trials, and the difference between the two. Average respiratory signals are overlayed for each condition (red lines). Dashed line represents inhalation onset. Black contours indicate statistical significance (permutation tests as described elsewhere, *P* < 0.05, FDR corrected). The source data for panel (B) is available in S4 Data.

### Low frequency phase modulates high frequency amplitude during inhale when odor is present

Previous work suggests that odor-induced theta oscillations predict odor identities, even during a detection task that does not require participants to identify odors [57]. Here, we found that theta rhythms in our data appeared to have a relationship with beta responses, which appeared to oscillate at around 5 Hz during inhale. Specifically, the overall magnitude of beta oscillations increased and decreased rhythmically with a frequency in the theta range, particularly in the first 1.5 to 2 seconds of the response (**Fig 6A**, gray box). Based on this observation, we hypothesized that theta oscillations might organize later-emerging higher frequency oscillations through phase modulation. To estimate modulation of higher frequency amplitude by theta phase, we calculated the modulation index (MI) [99] using theta phase as the modulating signal and a range of high frequencies (13–150 Hz) as the amplitude signals, for both odor and no-odor conditions (**Fig 6B**). We found that in the 1s time window following sniff onset, theta phase significantly modulated beta and gamma oscillations during the odor condition (*P* < 0.05, FDR corrected for MI in all frequency bands; peak MI at 16.16 Hz, mean z score across all frequencies = 7.52), but not during the no-odor condition (mean z score across all frequencies = 2.02).

**Fig 6.**
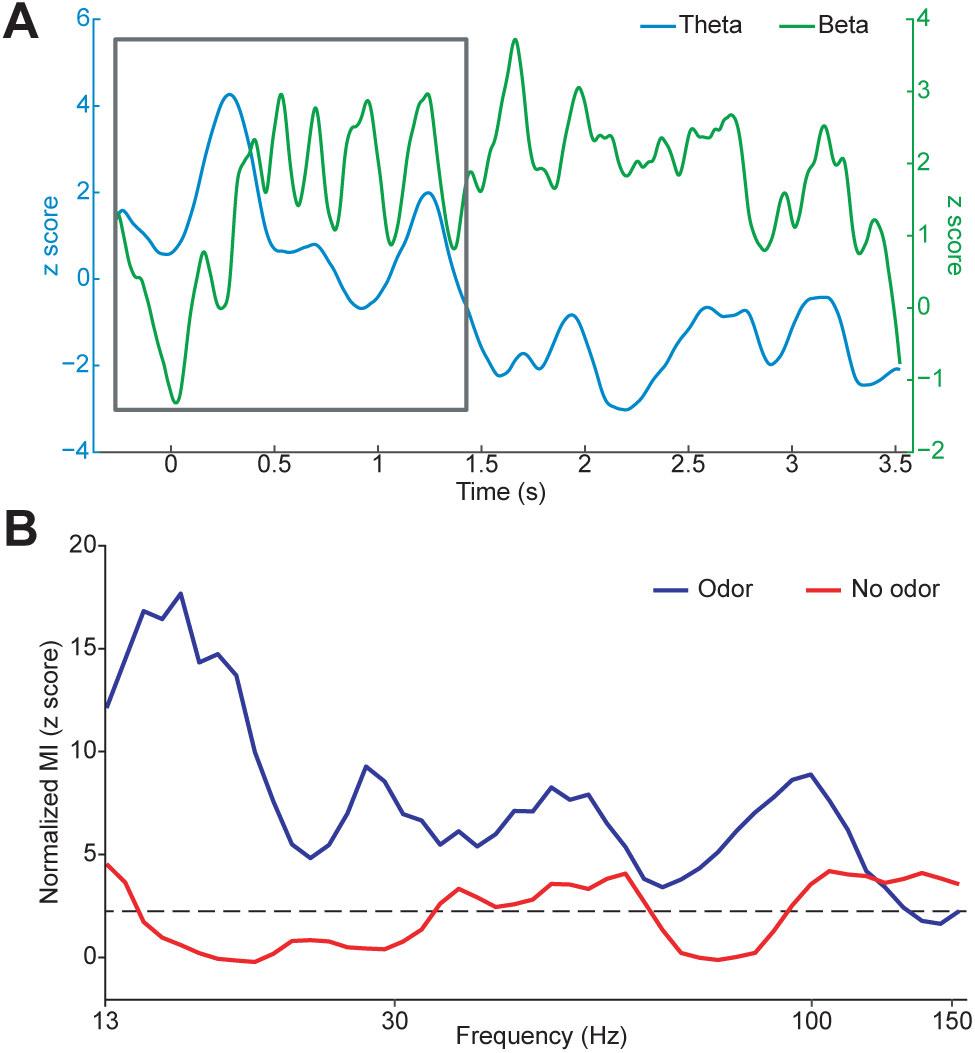
Theta phase modulates higher frequency amplitudes during inhale, when odor is present. (A) Apparent rhythmic beta amplitude modulations at the theta frequency. With effects of attention subtracted out, the average Z score of the amplitude of the odor responses over time is displayed for beta (green) and theta (blue) frequency bands. The overall magnitude of beta oscillations increased and decreased rhythmically with a frequency in the theta range, particularly in the first 1.5 to 2 seconds of the response (gray box). (B) Phase-amplitude coupling between theta phase and higher frequency amplitudes. Z-normalized modulation index (MI) of theta phase and higher frequency amplitudes (13 to 150 Hz), during 0–1s post sniff for all trials in the odor and no-odor conditions. Dashed line indicates FDR threshold for significant MI (*P* < 0.05, FDR corrected). The source data for panel (A) and (B) are available in S4 Data.

Our phase-amplitude coupling analyses was performed during a window in which a sensory-evoked potential could have occurred, and therefore the derived phase-amplitude coupling could have been driven by the evoked potential having a steep slope. To control for this possibility, we conducted a permutation analysis [100] in which we shuffled the trial-by-trial relationship between theta phase and higher frequency amplitude to test whether the modulation effect was due to the exact trial-by-trial relationship or rather induced by a steep slope during each trial. We normalized the observed MI to the trial-order-shuffled null distribution and still found significant effects for the odor condition but not the no-odor condition (**S5 Fig 5**). This finding suggests that higher frequency oscillations were modulated by theta phase during the early period of sniffing, when theta amplitudes were significantly increased.

### Theta, beta and gamma oscillations are present across a range of olfactory tasks

Studies in rodents suggest that different olfactory tasks may elicit distinct neural oscillatory signatures [32, 91]. Whether this is the case in the human olfactory system is unknown. It is therefore possible that the oscillations we observed in the theta, beta and gamma ranges were due to the nature of the olfactory task (identity matching to a cue, as opposed to detection [57] or pure identification). To explore this possibility, we present data from two additional subjects who performed 3 different olfactory tasks, including a detection task, an edibility assessment task and an odor naming task. This allowed us to determine if the nature of the task was the main driver of the effects we observed, rather than the odors. None of the tasks involved matching to a previous cue. For each task, trials began with a countdown to a cued sniff. Subjects self-initiated odor delivery using a manual hand-held olfactometer that allowed for precise, sniff-controlled timing of odor delivery. Following odor presentation, during the detection task, subjects indicated whether or not they detected an odor via button press. During the edibility assessment task, subjects indicated whether or not the odor was edible via button press. During the naming task, subjects indicated whether or not they could name the smell via button press, and then they spoke the name, if possible. All three of these tasks resulted in highly similar oscillatory responses to odor in both subjects (**Fig 7**). Both subjects, across all three task types, showed rapid increases in theta, followed by oscillatory increases in beta and gamma that persisted into exhalation. Detailed analyses of the responses to these different tasks is beyond the scope of this manuscript; results are displayed here solely for the purpose of this control analysis, which shows that the oscillations we observed in our original data set are present across a range of olfactory tasks, and were therefore unlikely to have been driven solely by the nature of the task.

**Fig 7.**
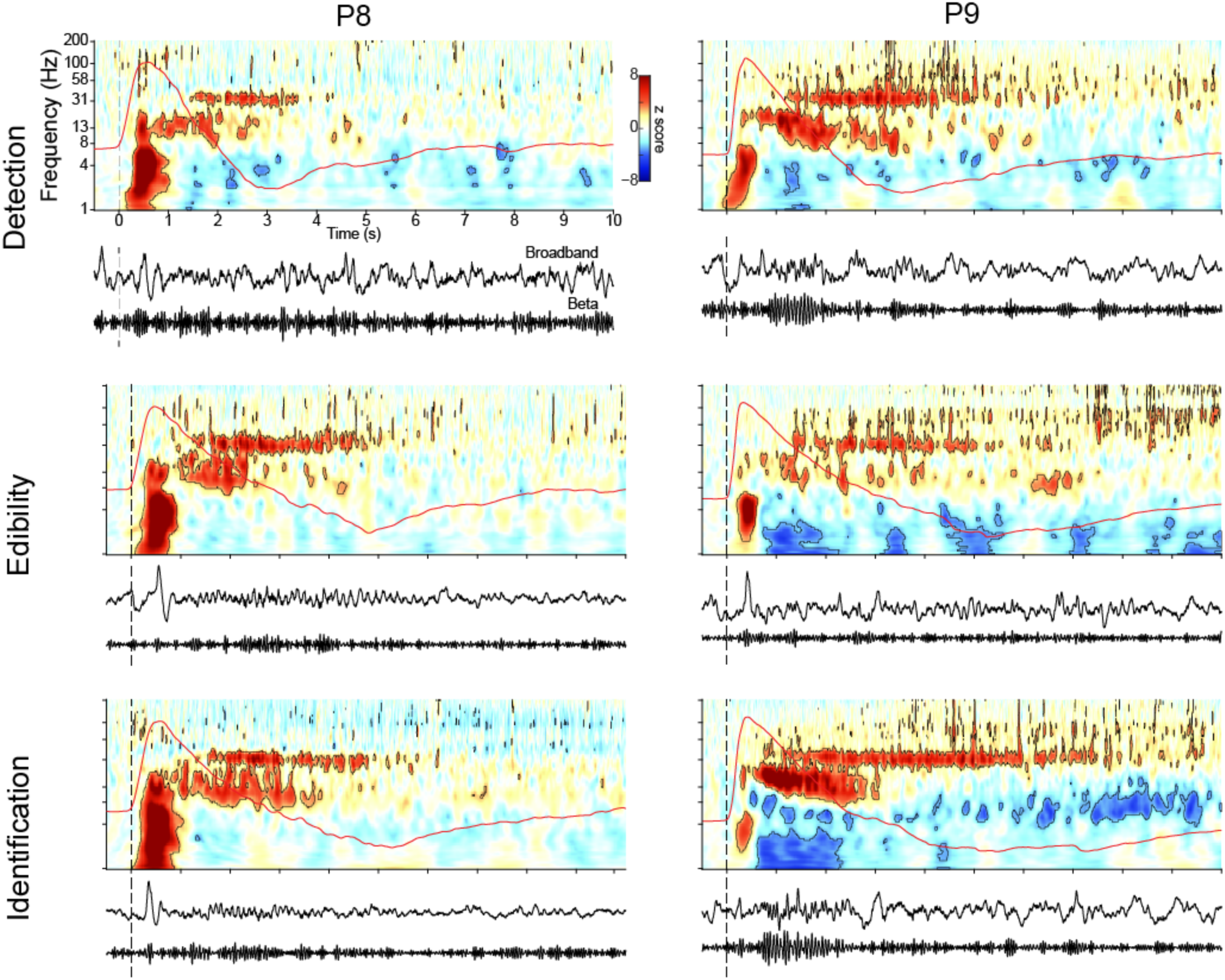
Odor-induced theta, beta and gamma oscillations are evident across a range of olfactory tasks. Shown are spectrograms from two participants (P8 and P9) who performed three different olfactory tasks including a detection task (top row), an edibility assessment task (middle row) and an odor naming task (bottom row). Short-dashed vertical line indicates sniff onsets. Average respiratory signals from each participant are shown as a red overlaid line on each spectrogram. Black contours indicate statistical significance (*P* < 0.05, FDR corrected). Below each panel is broad-band and beta-filtered time-series from one trial for that participant. The source data is available in S4 Data.

## Discussion

We found that odor induces oscillations in human piriform cortex in the theta, beta and gamma frequency bands, each with distinct temporal dynamics, forming the basis of a characteristic human olfactory cortical response. While theta oscillations rapidly emerged, peaked and dissipated within the first 500ms of inhalation, gamma and beta oscillations emerged and peaked later, extending into the exhale period (see **Fig 2**, and **Fig 3B and 3D**). These findings suggest that the spectrotemporal characteristics of oscillations in piriform cortex in response to odor are similar in rodents and humans. While our study cannot speak to the origin of these oscillations, it allows us to consider the possibility that knowledge of the drivers of LFP oscillations in rodents [8] may be applicable to humans, despite differences between species [101]; for example that beta and gamma oscillations in piriform cortex rely on an intact olfactory tract from the bulb [38], with important implications for our understanding of the top down or bottom up nature of olfactory processing [102–104] . We further found that the strength of beta and gamma oscillations was significantly more correlated with odor identification accuracy than theta oscillations, suggesting beta and gamma rhythms are important for odor identification in humans. Our work serves to unite results from previous intracranial human studies that have reported odor-induced oscillations at both low and high frequencies, with variable results [57,72–76,104].

Interestingly, we found that gamma rhythms were more tightly correlated to odor identification accuracy than beta during exhale (see Fig 4C). Since gamma rhythms are thought to reflect local computations [23,32,33,36,40,44,61,64], and beta oscillations are thought to reflect long-range network interactions [27,30,35,36,61,64], local computations in piriform cortex may have particular significance during exhale, though our study does not directly address this and future studies on this topic are needed. Our data also highlight an important role for theta oscillations in human piriform cortex, in agreement with previous studies [57], as we found that theta phase modulated beta and gamma amplitudes during inhalation, only when odor was present (see Fig 4). Rodent work has identified functionally distinct subpopulations of piriform neurons that differentially encode the intensity and identify of odor [82][81][105]. It is possible that the distinct oscillations we identified in human piriform cortex represent distinct populations of neurons that encode different features of the odor percept with unique temporal and spectral properties, though future work is required to determine this. In particular, the addition of single unit recordings may allow us to determine whether the different oscillations are tied to different subpopulations of piriform neurons.

Theta, beta and gamma oscillations have been extensively studied in the olfactory systems of rodents, providing insights into the functional roles of these oscillations in olfactory perception [23,24,35–39,41,43,60,64,65,26,68,106,107,27–31,33,34]. Furthermore, an impactful body of rodent work has shown that the timing of olfactory responses relative to sniffing behavior is an important mechanistic feature of odor coding, enabling accurate olfactory percepts to form quickly [33,41,116,108–115]. Though much of this work has focused on the olfactory bulb, responses in rodent piriform cortex also include temporal modulations [81,113,117], and its oscillations reflect sniff phase [26,62,78]. In contrast to the extensive and elegant body of rodent work, oscillations in human piriform cortex have been vastly understudied. Here, dovetailing with animal work, we found that theta, beta and gamma oscillations emerged at different times relative to sniff onset. The link between sniffing behavior and oscillations in rodents, combined with dramatic differences in sniffing behavior between rodents and humans, raises some intriguing questions about the timing of odor responses in humans. When sampling odors, rodents engage in multiple sniffing strategies, and may sniff at rates up to 12 Hz [118–120]. Humans might at times increase their sniffing frequency a bit when smelling an odor, but often sample odors in a single sniff, and aren’t able to reach sniffing frequencies of 12 Hz (though this has not yet been directly tested). Despite these large differences in sniffing behaviors across species, we found surprisingly similar time courses of odor responses in piriform cortex. For example, Frederick et al. [64] found that beta oscillations in rodent piriform cortex emerged at around 200 ms following odor onset, similar to our findings in human piriform cortex. We found that beta oscillations peaked and persisted longer than gamma oscillations, also in agreement with findings in rodents [59]. In rodents, the theta rhythm is actively shaped by sniffing behavior, which, in turn, shapes higher frequency oscillations. In humans, sniffing behavior almost never occurs at frequencies near the theta range, yet we found that theta oscillations shape higher frequency oscillations in the human olfactory system during inhalation, as is the case in rodents. This may suggest that the time scale of some odor-induced oscillations is not entirely dependent on breathing rate, and that network properties that support these rhythms may be conserved across species. Our findings support the idea that the theta rhythm serves as the “internal reference clock” of odor information processing [57], though future studies combining data from both humans and rodents, and potentially from noninvasive human olfactory bulb recordings [121] could contribute much-needed insights to answer this question.

An important consideration in comparing our findings to those in rodents is the nature of the task. The two most commonly used olfactory tasks in rodent studies are the Go-No Go (GNG) task and the 2-Alternative Forced Choice (2AFC) task [64,122–124]. The GNG task requires a response only when a target odor is present, whereas the 2AFC task requires a different response to each odor, depending on the identity of the odor. Our task is more similar to the 2AFC task, as it requires subjects to respond differently to each odor, depending on the identity of the odor. However, it is generally challenging to compare tasks between species—both GNG and AFC tasks require training of rodents, and include rewards during each trial. This is less common for human studies, in which experimenters directly instruct subjects on the task, and rewards are not required for subjects to learn the rules of the task. Several rodent studies have shown differences in oscillatory signatures across different olfactory tasks and difficulty levels [40, 64]. It is therefore possible that differences in task-related responses could impact differences between correct and incorrect trials between species [125–127]. A worthwhile direction for future research includes a direct comparison between rodent and human tasks, in which humans are trained and rewarded on a trial by trial basis, exactly as is typically conducted with rodents.

We found that the magnitude of beta and gamma oscillations strongly predicted odor identification accuracy, whereas this effect was weaker in the theta band, offering support for distinct functional roles for the different rhythms in olfactory processing [23,30,36,118]. Establishing beta rhythms in human piriform cortex may also lay the groundwork for future work on olfactory networks and integration of olfactory information with other cognitive processing streams and behaviors [30]. For example, integration of olfactory information with language networks [128–130], memory networks [131–135], sleep states [136–139], and other olfactory-guided behaviors [140, 141] could potentially involve interactions in beta oscillatory networks. Interestingly, we found that theta oscillations, but not beta or gamma oscillations, increased during attended states when no odor was present (**Fig 5B**), and prior to sniffing when odor was expected (**Fig 2A; Fig 3A and 3D**). Taken together, these findings may indicate a role for theta in olfactory attention, in line with previous studies suggesting that the phase and amplitude of lower frequency oscillations reflect attentional states in the human brain [97]. Olfactory attention also shapes neural responses in rodent olfactory cortex [95], and potential future work may include a direct comparison of these oscillations between species. It may also be interesting to explore these oscillations in other primary olfactory areas, which may have similar neural responses to piriform cortex.

A limitation of our work is that we were unable to fully tease apart the contributions of attention and expectation to our findings. Though we did conduct a control analysis designed to look for effects of attention and anticipation by analyzing responses to sniffs taken without anticipation of odor in between tasks, this analysis was limited by the fact that there was no odor during the sniffs that we analyzed. A perfect control would include presentation of odor in the absence of any expectation or anticipation. This would be difficult to achieve, as multiple presentations of odor across an experimental session almost undoubtedly would be anticipated after a couple of trials. This is an interesting topic for future studies, once these challenging aspects of experimental design are addressed.

Overall, our findings begin to define a characteristic human spectrotemporal odor response, consisting of three distinct frequency bands with unique time courses relative to sniff onset, with lower frequency phase modulating higher frequency amplitude, and corresponding closely with cortical responses that have been found in rodents. This lays the groundwork for future studies to further explore the functional role of the different aspects of the human cortical olfactory response.

## Methods

### Ethics statement

This study was approved by the Institutional Review Board at Northwestern University, under protocol #STU00201349, and the study adhered to the Declaration of Helsinki and the Belmont Report. Participation was voluntary and written informed consent was obtained from all participants.

### Participants

Our study included iEEG data from nine patients with medication-resistant epilepsy. All participants had depth electrodes implanted stereotactically for clinical pre-surgical evaluation (**Fig 1A**). Electrode locations were determined solely based on clinical need and, in our participant population, included piriform cortex within the medial temporal lobe. Data were acquired at Northwestern Memorial Hospital.

### Behavioral task

Participants performed a cued odor identification task in which they were asked whether an odor matched a prior auditory cue. This task involved periodic presentation of odors with inter-trial-intervals exceeding 6x the respiratory period of each individual, which was at least 15 s for every participant. During this inter-trial period, participants breathed naturally through the nose, and no odor was presented. The experimental trials were conducted in each participant’s hospital room, and were computer-controlled, presented to participants using an Apple laptop computer running MATLAB (RRID: SCR_001622) via the *PsychToolBox* extension (RRID: SCR_ 002881). Each trial of the task began with an auditory cue consisting of either the word “rose” or the word “mint”. The cue was delivered by computer speaker. After a delay of 3 to 7 seconds, the odor of rose (essential oil) or mint (methyl salicylate) was delivered through opaque squeeze bottles, while nasal airflow was monitored in order to precisely determine sniff—and therefore odor—onsets. We wanted participants to self-initiate odor delivery, so that they could control the timing of the stimulus, and to ensure high quality data by allowing patients to initiate trials at appropriate timing according to their individual situations. Note that this design increased the jittered timing between the cue and the sniff, with patients initiating the odor stimulus within 3 to 7 seconds following the cue. Participants were instructed that following the auditory cue, when they were ready, they should exhale, bring the bottle to their nose, and then sniff to sample the odor. For two patients, the experimenter held the bottle and followed the same procedure. At the moment when the patient sniffed, the experimenter sent a sync pulse to the clinical acquisition system via button press. The experimenter’s button press sent a signal via a data acquisition board (USB-1208F, Measurement Computing), which translated TTL pulses from MATLAB into the clinical EEG acquisition system (Nihon Kohden, Japan). Importantly, this button press did not provide sniff onset information—rather, it served to mark the odor-containing sniff in order to differentiate this sniff from other sniffs. Precise sniff onsets were determined by analysis of the nasal airflow signals, described in the methods. After smelling the odor, participants indicated whether the odor matched the cue via button press, with the exception of two patients who preferred to speak their response, which was recorded by an experimenter.

As previously described [98], participants completed between 48 and 64 trials, except for 1 participant who completed only 16 trials due to clinical constraints. The average inter-trial interval was 21.3 s, ranging from 14 to 28 s, across participants. The average performance on the task was 73.3% correct (P1: 79.69%, P2: 75.56%, P3: 31.25%, P4: 91.07%, P5: 100%, P6: 87.5%, P7: 46.88%), which means the response was “yes” when the odor matched with the sound cue or “no” when they were not matched. We found no difference in performance between the first (mean ± standard error: 73.98% ± 9.12%) and second (72.58% ± 10.72%) half of trials (two-tailed Wilcoxon signed-rank test; z = 0.40, *P* = 0.69). The performance was higher in cue-odor-matched trials (85% ± 7.08%) than nonmatched trials (58.17% ± 15.05%) (two-tailed Wilcoxon signed-rank test; z = 2.20, *P* = 0.028).

### iEEG and respiratory signal recording

iEEG signals were recorded using the clinical EEG data acquisition system (Nihon Kohden, Tokyo Japan) that is currently in use in Northwestern Memorial Hospital. The sampling rate for each participant was determined clinically, and ranged from 500 to 2000 Hz across participants. The reference and ground consisted of an implanted electrode strip on the surface of the brain facing the scalp. We recorded nasal airflow using a piezoelectric pressure sensor (Salter Labs Model #5500) with a nasal cannula placed at the patients’ nostrils during the experiment. Nasal airflow signals were recorded directly into the clinical acquisition system, and were therefore automatically synchronized with the iEEG data. The signal was first mean centered, then key respiratory features—including inhale and exhale onset, respiratory peak, and trough volume and time—were detected automatically with MATLAB toolbox *BreathMetrics*, developed by our lab [142], and then manually confirmed.

### Electrode Localization

As previously described [97, 98], to determine the implanted electrode locations, we used pre-operative structural MRI scans and post-operative computed tomography (CT) scans using the FMRIB Software Library’s (FSL) registration tool *flirt* [143, 144]. Individual CT images were registered to MRI images using 6 degrees of freedom and a cost function of mutual information, which was followed by an affine registration with 12 degrees of freedom. Individual MRIs were registered to a standard Montreal Neurological Institute (MNI) brain (MNI152_1mm_brain included in FSL) with 12 degrees of freedom. Finally, the transformation matrices generated above were combined to create a transformation from the individual CT image to standard MNI space.

The electrodes were localized by thresholding the raw CT image and calculating the un-weighted mass center of each electrode. Finally, the coordinates were converted to standard MNI space using the transformation matrix generated above.

Though we analyzed spectrograms from all electrodes on the PC depth wires and all electrodes on the parietal grids, electrodes corresponding to those shown in Fig 2 were selected by the following procedure. PC: For each subject, we first determined which subset of electrodes was anatomically located inside PC using pre-drawn individual ROIs based on the Human Brain Atlas [145]. This typically included between 1 and 3 electrodes. For the subjects who had only a single electrode in PC, we used that one. For subjects with multiple electrodes within PC, we chose the one that was closest to the center of PC. Notably, we also analyzed all piriform electrodes separately with similar results.

### Time-frequency analysis

For all time-frequency analyses, filtering was conducted using a two-pass, zero-phase-lag, finite impulse response (FIR) filter, as implemented by the MATLAB toolbox *fieldtrip* (RRID: SCR_004849), unless specified otherwise. We first low-pass filtered the iEEG signal at 235 Hz and then removed 60 Hz noise and its harmonics with a band-stop filter with a bandwidth of 4 Hz. We then downsampled the signal to 500Hz and re-referenced the data to the common average. To compute spectrograms, we filtered the pre-processed data between 1 to 200 Hz in 100 logarithmically increasing steps ranging from 2 to 50 Hz in width. We kept only the first 95 frequency bands from 1 to 153 Hz as our frequencies of interest for all subsequent analyses unless specified otherwise. The analytical amplitudes of the filtered signals were calculated by taking the absolute value of the Hilbert-transformed signal, and temporally smoothed with a moving average filter kernel of 10 ms.

To compute spectrograms, we first created sniff-onset-aligned epochs, extending from – 2 s prior to 4 s following the sniff onset, for each trial. Sniff onsets were determined using *BreathMetrics* [142]. Sniffs taken during experimental trials at the time of odor presentation were used for the odor condition, sniffs taken in between trials when no odor was present were used for the no-odor condition, and sniffs taken during experimental trials in between the cue and odor were used for the attended-no-odor condition. There were a larger number of no-odor trials compared to odor trials, and therefore, no-odor trials were randomly sampled without replacement for each participant separately to match trial numbers across conditions for further analysis. The spectrograms were calculated by averaging the amplitude epochs across trials at each frequency, which was further normalized by subtracting the baseline average. The time window of [−0.55, −0.05] s prior to cue onset was used as baseline. To determine statistical significance, we used a permutation method [97,98,146] to generate z score values. For each repetition of the permutation process, the sniff onsets were circularly shifted in time by a random amount while maintaining the relative distance between them. Then, we calculate he average amplitude across these randomly shifted events. The procedure above was repeated 10,000 times resulting a null distribution of baseline amplitude. Finally, the baseline-corrected real spectrogram was divided by the standard deviation of this null distribution, resulting a z score map. Raw power, baseline-corrected power and z normalized maps can be seen in **S6 Fig 6**.

To compare the spectrograms between odor and no-odor condition, we used a permutation method. In each permutation, the condition labels were shuffled across all trials, and the difference of baseline-corrected spectrogram between permuted odor- and no-odor-conditions were calculated. This process was repeated 10,000 times resulting in a null distribution of spectrogram difference at each time-frequency point. The mean and stand deviation of the null distribution was obtained using MATLAB’s *normfit* function. Finally, a z score of the real spectrogram difference was calculated by subtracting the mean and then dividing by the stand deviation. The spectrograms created for the attentional control analysis shown in Fig 4B were generated with the same methods, with the number of trials adjusted to include equal numbers of odor and attended-no-odor trials.

To compute the percentage change of the amplitude time series shown in **Fig 3A**, we averaged the amplitude time series across frequencies within each of the following frequency bands: theta 4–8 Hz, beta 13–30 Hz, and gamma 30–150 Hz, for each trial. The percentage change of amplitude was calculated as (Amplitude-Amplitude_baseline)/Amplitude_baseline for each trial. The baseline window was defined as −500 ms to −50 ms relative to cue onset. To compare the percentage change of amplitude between odor and no-odor conditions, we performed a paired sample t test at each time point. Multiple comparisons across all time points were corrected using FDR method.

In order to visualize the differences in temporal dynamics across frequency bands, we created a z-normalized amplitude map for each frequency band using a bootstrapping method (**Fig 3B**). First, each trial was baseline corrected and z-normalized. Then, in each repetition, we resampled all 323 trials (with replacement), and calculated the average across trials for each frequency band. A z-score matrix (repetition x time) was generated for each frequency band after 2000 bootstraps (**Fig 3B**).

We then performed a cluster-based analysis to quantify the time course of significant responses for each frequency band (**Fig 3C**). First, we performed a one sample t tests against zero for z-normalized amplitudes for all trials at each time point for each frequency band. To identify continuous significant clusters, We then used a cluster-based statistical thresholding analysis to identify continuous significant clusters. The *P* value threshold for initial clustering was set to *P* = 0.01, and the cluster size was defined as the sum of the t-statistics of a given cluster. To generate a null distribution of the cluster size, we used a permutation method. In each repetition (10000 repetition in total), a random number of trials were circularly shifted separately, and the maximal cluster size was calculated as described above. The 95^th^ percentile of the distribution was used as the cluster size significance threshold, which corresponds to cluster-based corrected *P* < 0.05.

We next conducted an analysis to explore the timing and phase of responses in each frequency band (**Fig 3D**). we calculated the distribution of respiratory phase values at which the peak amplitude responses occurred using a bootstrapping method. For each bootstrap, we resampled 323 trials (with replacement) while keeping the pairwise relationship between LFP and breathing signals. The LFP and respiratory signals were first averaged over resampled trials. To find the peaks of the LFP amplitude response, we smoothed the amplitude time series using a moving average method with a kernel of 50 ms. Then, the respiratory phase of the maximal amplitude over the 4 s post-sniff time period was extracted. The respiratory phase was obtained from the breathing signal using Hilbert method. Finally, the distribution of the respiratory phases across trials were qualified using PLV [147] as implemented in the MATLAB toolbox circstats (RRID: SCR_016651) [148]. The Rayleigh test was performed to test for the non-uniformity of phase distributions. For the peak timing analysis on the single participant level, we used the same method.

We next conducted a series of analyses to look for relationships between LFPs and odor identification accuracy (**Fig 4**). First, we computed amplitude spectrogram for odor and no-odor conditions separately. The two spectrograms were z-normalized to the same combined null distribution generated using methods described in the previous section to rule out the effect of trial number difference across conditions. In order to statistically compare the amplitude difference between correct and incorrect trials, we conducted a resampling analysis. For each repetition, we resampled equal numbers of correct and incorrect trials (71, with replacement) to ensure a fair comparison across conditions. On each repetition, we calculated the mean of z-normalized amplitudes across the entire time window for each frequency band for each condition (these distributions are shown in **Fig 4B**), subtracted the mean of incorrect trials from the mean of correct trials, and repeated this process 200 times to generate the distribution of difference values between correct and incorrect trials. The difference values for each frequency band was tested against 0 using one-sample t tests to generate statistics. The distribution was plotted as scatter as well as box plot (**Fig 4B**).

We then conducted a bootstrap-correlation analysis to directly assess the correlation between task performance and LFP amplitude of each frequency band for inhale and exhale respectively on a trial-by-trial basis (**Fig 4C**). For each repetition (1000x) in the analysis, we resampled 323 trials with replacement and calculated the accuracy of the chosen trial set. We then calculated the mean amplitude during inhale and exhale separately (inhale-exhale transition defined by phase angle of pi/2 for averaged breathing signal) for all three frequency bands. We calculated Pearson’s correlation coefficients between task accuracy and mean amplitudes for inhale and exhale, for each frequency band. We then repeated this entire process 50 times to generate a distribution of correlation coefficient for each preparatory phase and frequency condition. We used FDR to correct for multiple comparisons (*P* < 0.05) when we set the significance threshold for the r value. We subsequently fisher-z-transformed all Pearson’s r values and conducted a repeated measures two-way ANOVA as well as paired sample t tests to analyze the differences across frequency bands and respiratory phases.

We next created scatter plot visualizations of amplitude responses during inhale and exhale phases for correct and incorrect trials and fitted a linear SVM classifier to each of the scatter plot in order to see how well the amplitude during inhale and exhale, taken together, could predict task performance **(Fig 4D** and **4E).** To balance the discrepancy between correct and incorrect trial numbers (252 vs. 71), we conducted a bootstrapping analysis using equal numbers of trials across conditions on each repetition. For each respiratory phase and frequency band, we sampled 30 trials within condition with replacement each time, calculated percent change relative to baseline with the trial-wise averaged amplitude time series, and calculated average amplitudes during inhale and exhale for each trial set. Inhale and exhale periods were defined as 0–1.5s and 1.5–3 s post sniff respectively for all trials. This process was repeated 500 times for both correct and incorrect trials to generate a scatterplot of corresponding inhale and exhale amplitudes for trials from both conditions. In order to test the separability of data, we applied a linear binary SVM classifier using MATLAB function *fitcsvm* with five-fold cross validation. ROC curves were generated using the function *perfcurve*.

### Phase-amplitude coupling analysis

We calculated the modulation index (MI) to measure coupling between theta phase and higher frequency amplitudes (13–150 Hz) (**Fig 5**). To do so, we extracted the phase angle time series for the theta frequency band (4–8 Hz), and then from that phase time series, created sniff-aligned trial epochs. Data from all trials was concatenated. To generate the higher frequency amplitude data that was used to compute the MI, the raw time series were filtered and amplitude-extracted (as described previously) to the same 47 log-spaced frequency bands (13.05–153.04 Hz), with logarithmically increasing bandwidth, as described in the previous section for the time-frequency analysis. We then concatenated this data identically to the theta phase data. MI was calculated as the Kullbach-Leibner distance divided by the logged number of bins. To normalize the MI, we created a null distribution of MI values using permutation method. For each repetition, and for each frequency band, we circularly shifted the amplitude time series, and calculated the MI between the shifted amplitudes and theta phase values. This process was repeated 1000 times for each frequency band to generate the corresponding null distribution of MI values. We then normalized the raw MI to the null distributions to generate the z-scores, and FDR corrected the *P* values for multiple comparisons. To control for the effect of evoked potential on phase-amplitude coupling, we used a permutation method to ensure that the significant MI we observed was unlikely due to steep slope of evoked potential occurring in every trial. For each repetition, we shuffled the order of theta phase but kept the order of higher frequency amplitude to randomize the trial-by-trial relationship, and calculated MI for each frequency band based on the shuffled trials. This process was repeated 1000 times for each frequency band to create a null distribution for the MI value under the null hypothesis that the phase-amplitude coupling was due to steep waveform occurring in every trial. We then normalized the raw MI to the null distributions to generate the z-scores, and FDR corrected the *P* values for multiple comparisons.

### Control for nature of the task

Two participants performed three different olfactory tasks over six separate blocks. Patients performed two blocks of each task. The three tasks required the participant to indicate: 1) whether or not they detected any odor (Detection task), 2) whether the odor was edible (Edibility task), or 3) the identity of the odor (Naming task). For all three tasks, each trial began with a countdown (“3, 2, 1, Sniff”) which appeared in sequential order, with each number lasting 1 s. The participant was instructed to sniff when the word sniff appeared, using a manual hand-held olfactometer that allowed for sniff-controlled timing of odor delivery. The hand-held olfactometer consisted of an odor canister connected to a nose port, constructed with one-way valves at the outlet and inlet such that the headspace was able to be sampled only and immediately upon sniffing. The canister was contained inside of an opaque 3D printed case. Each run consisted of 20 trials, resulting in 40 trials of each task type. Odors were of natural origin, and included banana, salsa, liquid smoke, blood orange tea, peanut butter, garlic, peppermint, pine and rose, for each trial. The tasks were performed while clinical ongoing electrophysiological activity was recorded at a sampling rate of 1000 Hz using a 256-channel clinical EEG acquisition system (Nihon Kohden). The respiratory signals were recorded using a pressure sensor at the nose (Salter Labs). Time-frequency analysis of odor-induced responses in the piriform cortex was conducted using the methods described above.

## ACKNOWLEDGEMENTS

We thank the patients and families of patients who volunteered to participate in this research; without their generous contributions of time and effort, this study would not have been possible. We thank Navid Shadlou for his technical expertise and support, and assistance with data collection. This work was supported by National Institutes of Health grants R01-DC016364 (NIDCD) and R01-DC018539 (NIDCD) to C. Z.

---

**S1 Fig.**
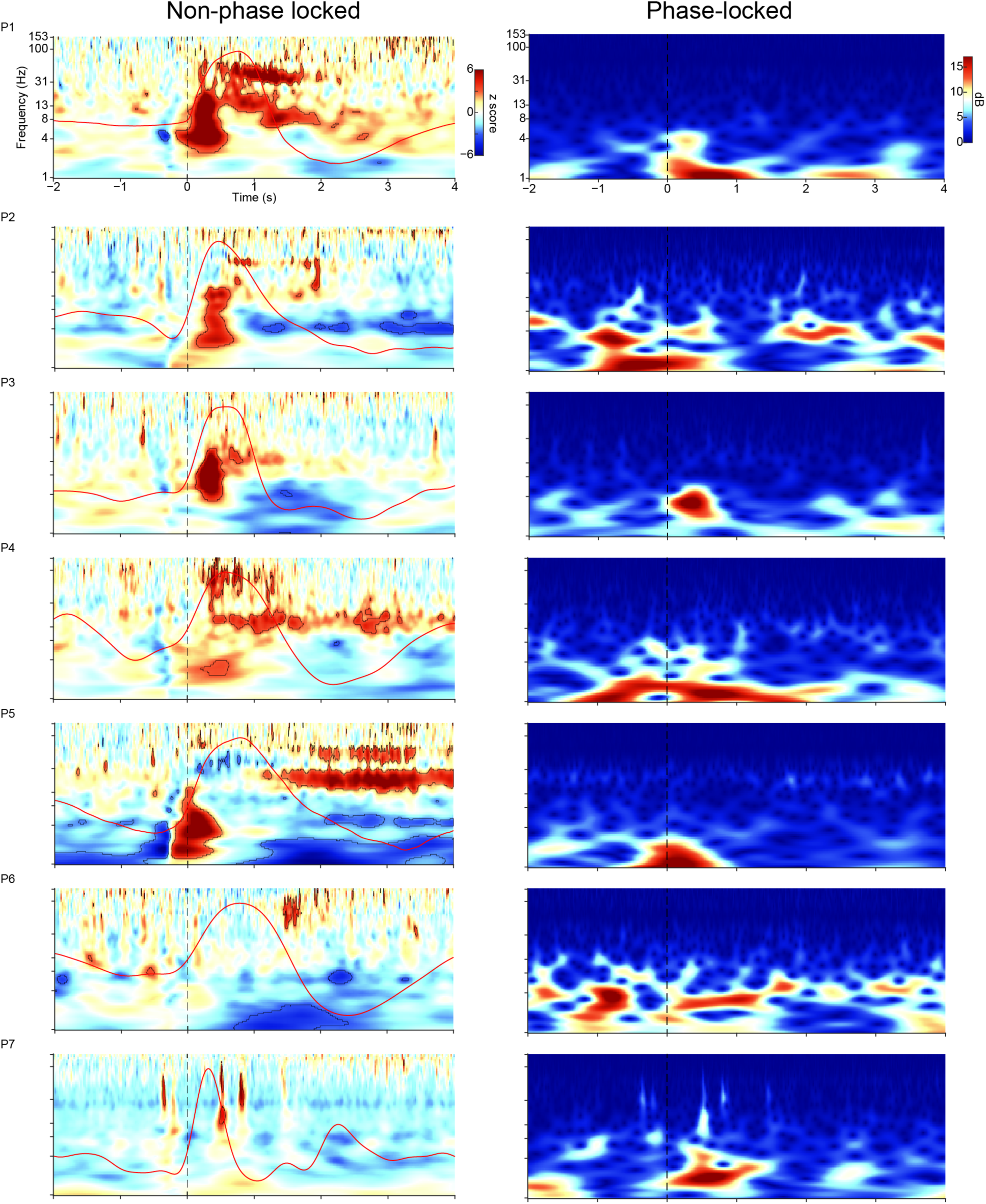
Related to Fig 2. Odor-induced and odor-evoked spectrograms. Non-phase-locked (left) and phase-locked (right) spectrograms are shown for each participant (P1–P7). The non-phase-locked spectrogram was obtained by subtracting the event-related potential from each trial before time-frequency decomposition. The red solid overlay indicates each participant’s respiratory signal. Black outlines indicate statistically significant clusters (*P* < 0.05, false-discovery-rate corrected). The phase-locked spectrogram was calculated as the baseline-corrected time-frequency decomposition of the event-related potential. The vertical short-dashed lines indicate sniff onset. The source is available in S5 Data.

**S2 Fig.**
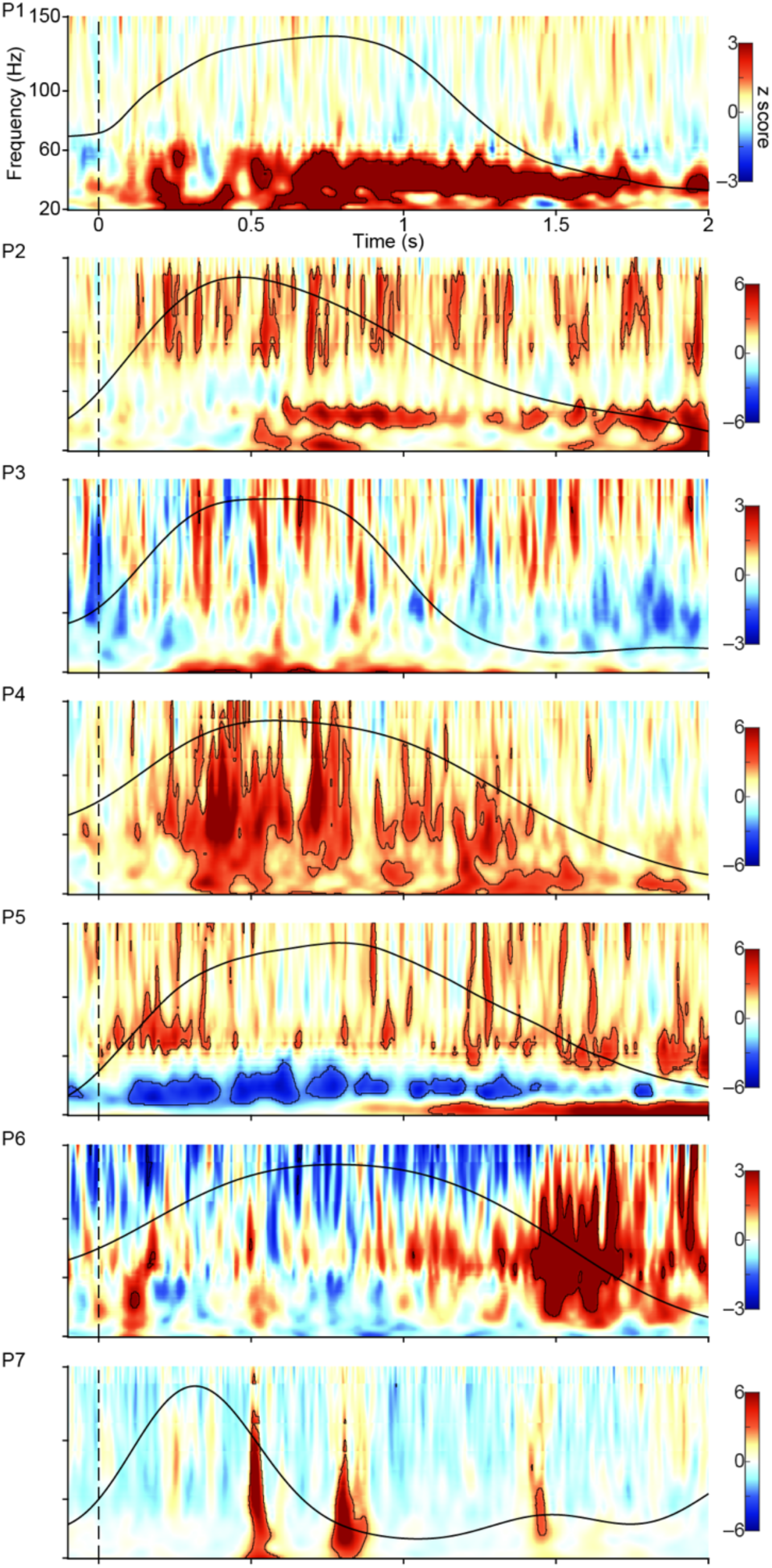
Related to Fig 2. Sniff-onset aligned spectrograms with linear frequency scale. The black solid overlay indicates each participant’s (P1–P7) respiratory signal. Black outlines indicate statistically significant clusters (P < 0.05, false-discovery-rate corrected). The vertical short-dashed line indicates sniff onset. The source is available in S5 Data.

**S3 Fig.**
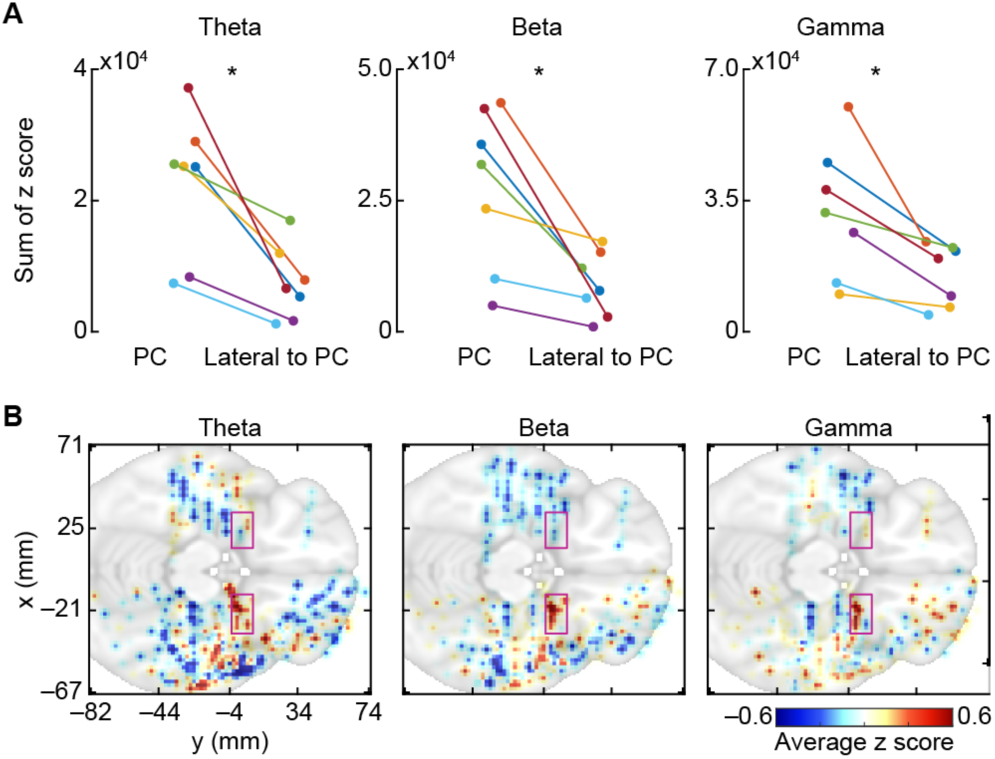
Related to Fig 2. Odor-induced responses are maximal in piriform cortex. (A) Odor-induced amplitude (sum of z score) is larger in the depth wire located in the piriform cortex (PC) compared to those located outside of the piriform cortex (Lateral to PC) in theta, beta and gamma frequency bands. * indicates statistically significant difference (two-tailed paired t test). (B) The mean z score was calculated over a time window of 2s in the theta, beta or gamma frequency band for each electrode and each participant. The data were collapsed over the z-axis and smoothed. The background brain is a slice (z = –16) of the Montreal Neurological Institute standard brain. The pink rectangles outline piriform cortex. The source is available in S5 Data.

**S4 Fig.**
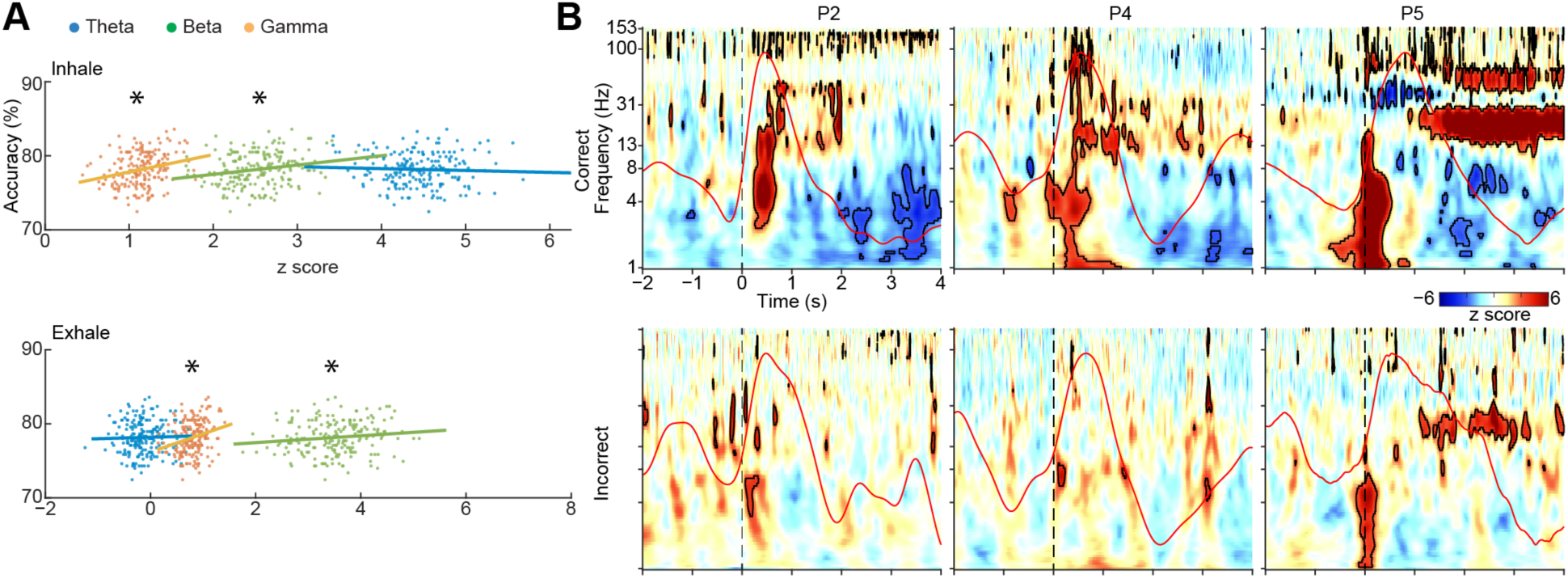
Related to Fig 4. Odor-induced responses for correct and incorrect trials. (A) Example scatter plots showing the correlations plotted in Fig 4C. The scatter plot corresponding to the correlation value of one dot from each bar in Fig 4C is shown. (B) Representative spectrograms from 3 participants, showing correct (top) and incorrect trials (bottom) separately. The red solid overlay indicates each participant’s (P1–P7) respiratory signal. Black outlines indicate statistically significant clusters (P < 0.05, false-discovery-rate corrected). The vertical short-dashed line indicates sniff onset. The source is available in S5 Data.

**S5 Fig.**
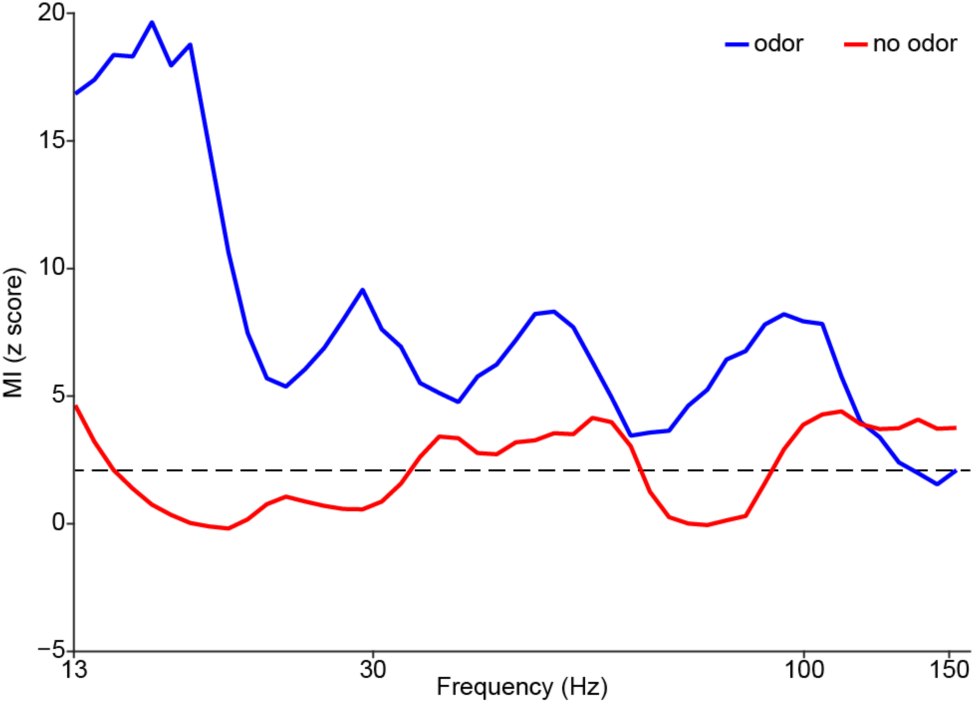
Related to Fig 6. Results of modulation index computed for odor and no-odor trials, accounting for possible contribution of steep slope of sensory-evoked potentials (see supplementary Methods). The source is available in S5 Data.

**S6 Fig.**
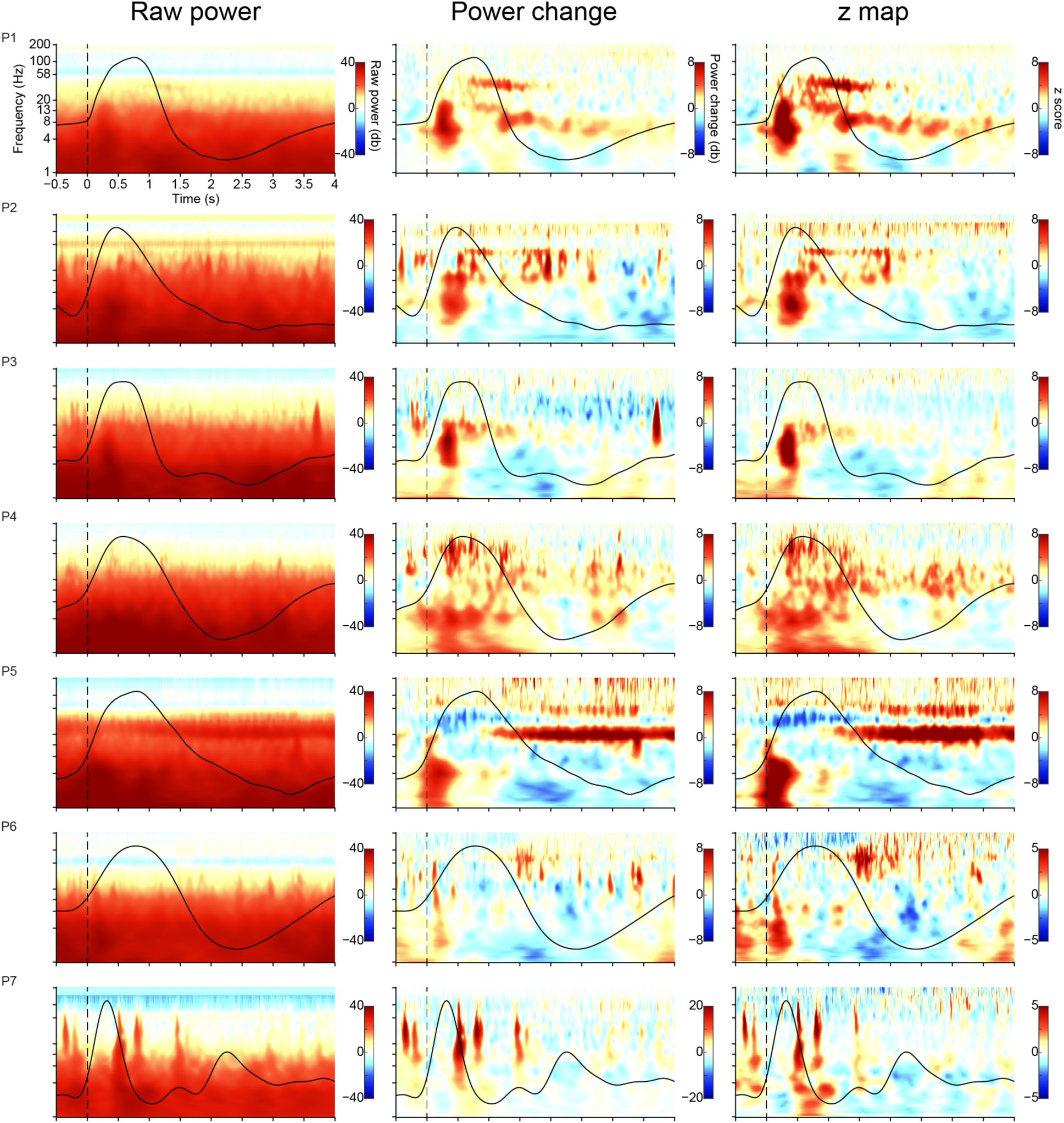
Related to Fig 2. Sniff-onset aligned raw power (left), power change relative to baseline (middle), and z score map (right, same as Fig 2C) are shown for each participant (P1–P7). The baseline was defined as [–0.55, –0.05] s prior to cue onset. The black overlaid line indicates the respiratory signal. The vertical short-dashed line indicates sniff onset. The source is available in S5 Data.

S1 Text. Supplementary methods.

S1 Data. Zip file containing datasets underlying Fig 2A, 2B, and 2C.

S2 Data. Zip file containing datasets underlying Fig 3A, 3B, 3C, and 3D.

S3 Data. Zip file containing datasets underlying Fig 4A, 4B, 4C, 4D, and 4E.

S4 Data. Zip file containing datasets underlying Figs 5B, 6A, 6B, and 7.

S5 Data. Zip file containing datasets underlying S1 Fig, S2 Fig, S3 Fig, S4 Fig, S5 Fig and S6 Fig.

